# Neolithic genomes reveal a distinct ancient HLA allele pool and population transformation in Europe

**DOI:** 10.1101/851188

**Authors:** Alexander Immel, Christoph Rinne, John Meadows, Rodrigo Barquera, András Szolek, Federica Pierini, Julian Susat, Lisa Böhme, Janina Dose, Joanna Bonczarowska, Clara Drummer, Katharina Fuchs, David Ellinghaus, Jan Christian Kässens, Martin Furholt, Oliver Kohlbacher, Sabine Schade-Lindig, Iain Mathieson, Andre Franke, Johannes Krause, Johannes Müller, Tobias L. Lenz, Almut Nebel, Ben Krause-Kyora

## Abstract

The Wartberg culture (WBC, 3,500-2,800 BCE) dates to the Late Neolithic period, a time of important demographic and cultural transformations in western Europe. We perform a genome-wide analysis of 42 individuals who were interred in a WBC collective burial in Niedertiefenbach, Germany (3,300-3,200 cal. BCE). Our results highlight that the Niedertiefenbach population indeed emerged at the beginning of the WBC. This farming community was genetically heterogeneous and carried a surprisingly large hunter-gatherer ancestry component (40%). We detect considerable differences in the human leukocyte antigen gene pool between contemporary Europeans and the Niedertiefenbach individuals whose immune response was primarily geared towards defending viral infections.

## Introduction

Over the last few years, large-scale ancient DNA (aDNA) studies have provided unprecedented insights into the peopling of Europe and the complex genetic history of its past and present-day inhabitants^1 2,3 4,5^. Recent research has particularly focused on the population dynamics during the Neolithic period. The first agriculturalists across central Europe, who are associated with the uniform Linear Pottery culture (Linearbandkeramik, LBK, 5,450-4,900 BCE) across central Europe, probably co-existed with local hunter-gatherers (HG) for about two thousand years^6^. Although these groups are thought to have lived in close proximity, initially only limited admixture occurred^2,3^. This situation changed later (4,400-2,800 BCE) when the gene-pool of the early farmers was transformed through the introgression of genomic components typical of HG populations^1,3,7^.

The Late Neolithic period is archaeologically characterized by strong regional diversification and a patchwork of small units of classification (i.e. archaeological cultures)^8^. One of the western units that emerged at the beginning of the Late Neolithic period is associated with the Wartberg culture (WBC, 3,500-2,800 BCE), which most likely developed from the Late Michelsberg culture (MC, 3,800-3,500 BCE)^9,10^. WBC is mainly found in western central Germany (Fig. 1)^11,12^. It is known for its megalithic architecture of large gallery graves that is distinct from that in adjacent regions, but shows a striking resemblance to similar monuments in the Paris Basin and Brittany^13,14^. Despite the central geographical location of WBC that connects cultural influences from several directions, no genomic data of human remains from WBC sites have so far been investigated.

**Figure 1:**
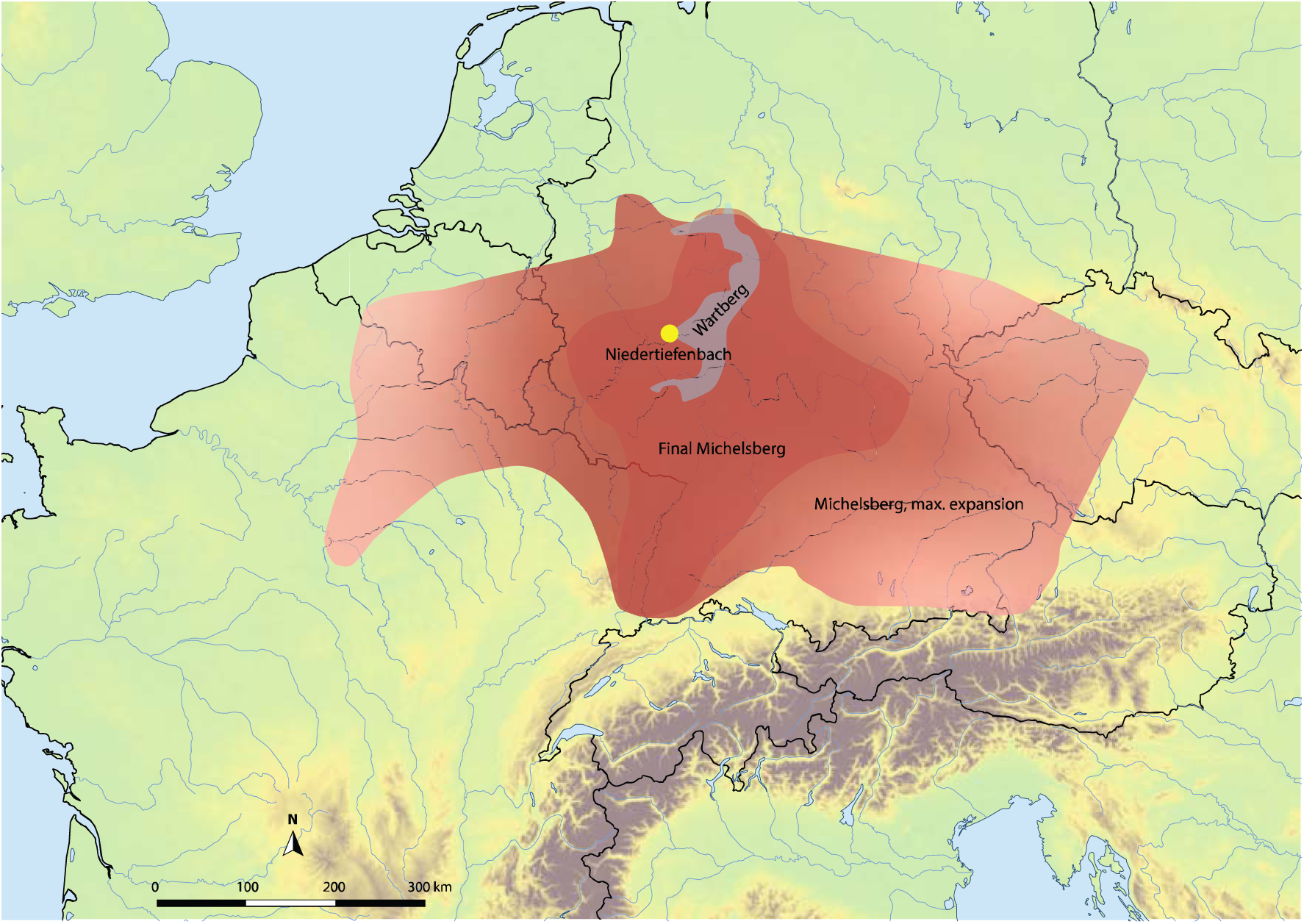
Map with the site Niedertiefenbach from where the individuals presented in this study were recovered. The temporal and geographic distributions of the archaeological small units mentioned in this study are shown.

Here, we performed a genome-wide data analysis of 42 individuals who were buried in a WBC gallery grave near the township of Niedertiefenbach in Hesse, Germany (Fig. 1), dated between 3,300-3,200 cal. BCE (Supplementary Information). In contrast to other genome-wide aDNA studies, which usually include a small number of individuals from a specific site and period, we provided a snapshot of a burial community that used the collective grave for approximately 100 years^15^ (Supplementary Information). In addition to population genetic and kinship analyses, we also investigated the human leukocyte antigen (HLA) region. This approach allowed us not only to reconstruct the genetic ancestry of the WBC-associated people from Niedertiefenbach, but also to gain insights into the makeup of immunity-related genes of a Late Neolithic group.

## Results

In total, aDNA extracts obtained from 89 randomly selected individuals interred in the Niedertiefenbach grave^16, 17^ were subjected to shotgun sequencing. Of these, we filtered out 47 who i) had fewer than 10,000 single-nucleotide polymorphisms (SNPs) benchmarked on a previously published dataset of 1,233,013 SNPs^1,2,4^ or ii) showed evidence of X-chromosomal contamination (>=5%). Thus, after quality control, datasets from 42 individuals were available for subsequent comprehensive analyses (Supplementary Table 1). aDNA damage patterns^18^ were consistent with an ancient origin of the isolated DNA fragments. When we screened the sequence data for known blood-borne pathogens such as *Yersinia pestis, Mycobacterium tuberculosis* and *Mycobacterium leprae* with MALT^19^, no signs of an infection were detected. Ten individuals were genetically determined as females and 25 as males. Although of the remaining seven individuals one was osteologically identified as female and one as male.

First, we applied principal component analysis (PCA) to project the SNP information derived from the Niedertiefenbach collective together with previously published datasets of 122 ancient populations onto a basemap calculated from 59 modern-day West-Eurasian populations^1,2,4,20^. The Niedertiefenbach individuals formed a cluster that is mainly explained by genetic variation between HG and early farmers on the first principal component (Fig. 2). However, the Niedertiefenbach samples covered a wide genetic space which reflects a high intra-population diversity. Some of the individuals grouped closely with those from the Blätterhöhle, a cave site near Hagen, Germany (4,100-3,000 BCE)^6^ (Fig. 2). ADMIXTURE analysis^21^ with four to eight components suggested two main genetic contributions to the Niedertiefenbach collective – one maximized in European HG and the other in Neolithic farmers from Anatolia (Fig. 3; Supplementary Fig. 1). Next we applied *f3* outgroup statistics^22^ to calculate the amount of shared genetic drift between the Niedertiefenbach sample and another test population relative to an outgroup [*f*_*3*_(*Niedertiefenbach; test; Mbuti*)]. The highest amount of shared genetic drift was observed between Niedertiefenbach and European HG from Sicily, Croatia, and Hungary (Supplementary Fig. 2). To estimate the amount of Neolithic farmer and HG genetic ancestry in the Niedertiefenbach group, we ran qpADM^22^. We obtained feasible models for Niedertiefenbach as a two-way mixture of Neolithic farmers from Anatolia and various European HG, which altogether gave on average ∼60% farmer and ∼40% HG ancestry (Supplementary Table 2). Another feasible two-way admixture model for Niedertiefenbach was as the combination of Anatolian Neolithic farmers (41%) and individuals from the Blätterhöhle. We then applied ALDER^23^ to estimate the date of admixture. We observed significant results for a mixture of components associated with early farmers and Loschbour HG (Waldbillig, Luxembourg) in the Niedertiefenbach population 14.85 +/-2.82 generations before the ^14^C benchmark of 3,300-3,200 cal. BCE (Supplementary Note 1). Based on a generation time of 29 years^24^, the date for the emergence of the genetic composition of the Niedertiefenbach community appears to be between 3,850-3,520 cal. BCE. However, the dates are based on the idealized model of a single wave of admixture between Anatolian Neolithic farmers and Loschbour HG. These populations were used as closest unadmixed genetic proxies for possible parental sources based on the qpADM results. The models do not take into consideration multiples waves, continuous admixture or admixture of populations that were already admixed^3^. Thus, the obtained dates reflect only the minimal number of generations.

**Figure 2:**
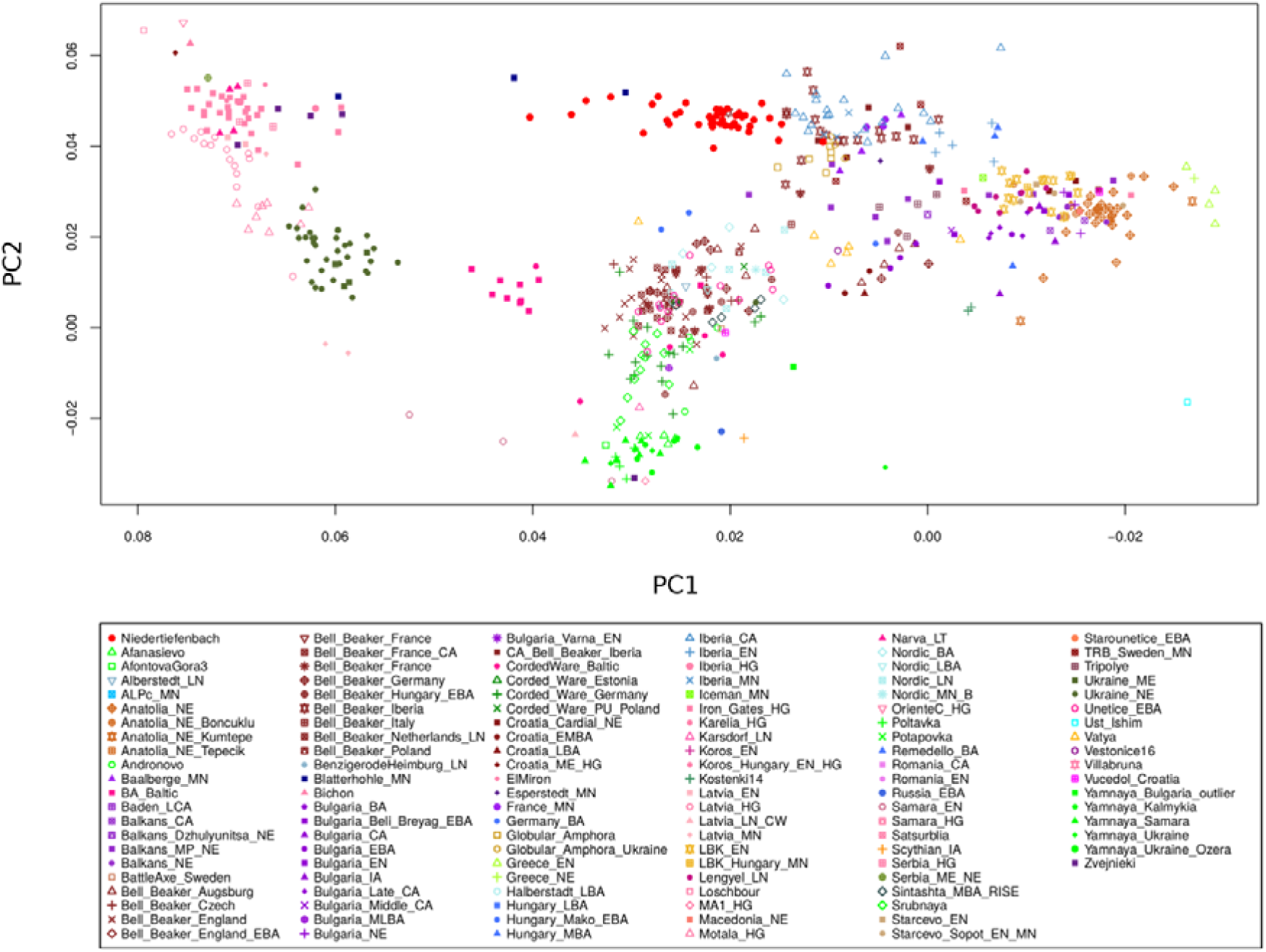
PCA of individuals from 123 ancient populations including Niedertiefenbach projected onto the first two principal components calculated from 59 present-day West-Eurasian populations (not shown for clarity). Niedertiefenbach individuals are depicted as red dots.

**Figure 3:**
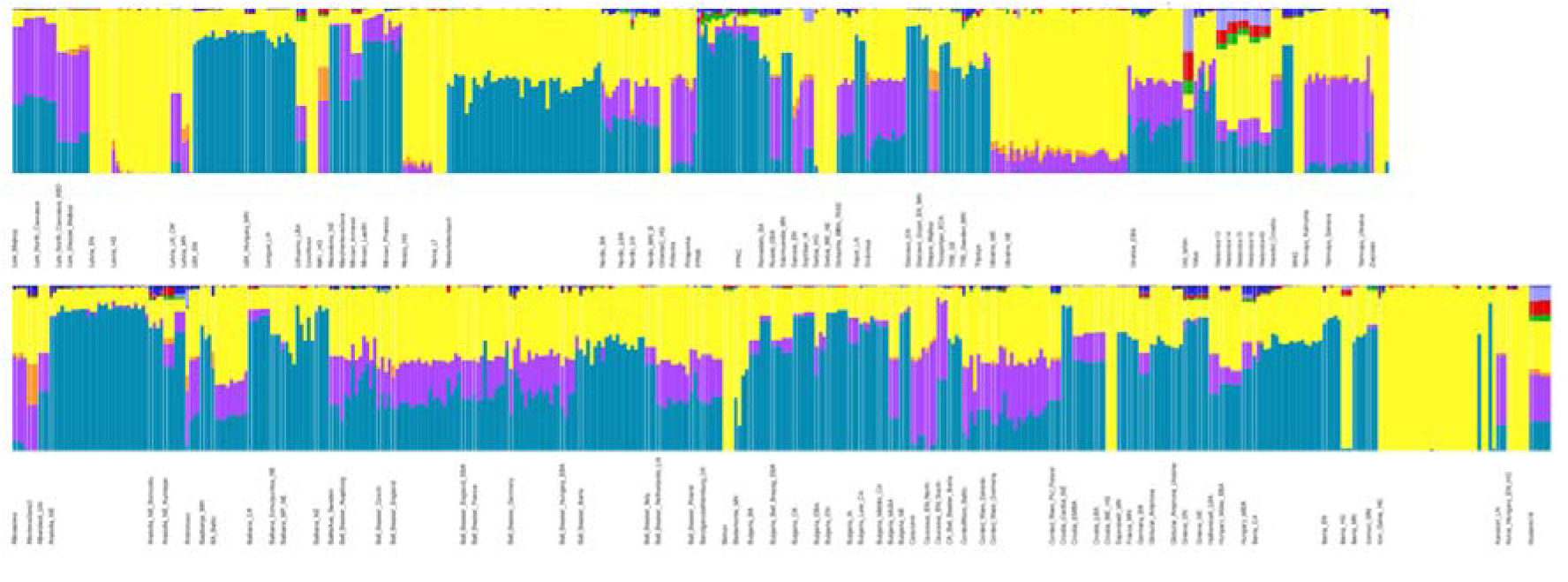
ADMIXTURE modeling of ancient populations including Niedertiefenbach with K=8 genetic components.

For phenotype reconstruction, we investigated selected SNPs associated with skin pigmentation and hair color (rs16891982), eye color (rs12913832), starch digestion (rs11185098) and lactose tolerance (rs4988235)^25^. Not all of these SNPs were available for all of the investigated individuals due to poor sequence coverage. Fourteen of the 42 individuals carried only the rs16891982-C allele, which is associated with dark hair and increased skin pigmentation^26^, while three had both alleles (C and G). Only three individuals carried the rs12913832-G allele associated with blue eye colour, seven had the A allele associated with brown eye colour, and eight had both alleles. The minor A allele of rs11185098 is positively associated with AMY1 (amylase 1) gene copies and high amylase activity, responsible for starch digestion^27^. Only one individual was found to be homozygous for the G allele and six had both alleles, while no homozygous carrier for the A allele was found. Interestingly, all individuals with enough coverage for the rs4988235 SNP carried the G-allele that tags an ancestral haplotype associated with lactose intolerance^28^, which suggests that the Niedertiefenbach people could not digest dairy products.

To determine the HLA class I and II alleles of the Niedertiefenbach individuals, we applied a previously developed method^29^. In addition, we used OptiType, an automated HLA-typing tool^30^. Only alleles that were consistently called by both methods were considered for the analysis. We successfully genotyped HLA A, B, C, DPB1, DQB 1 and DRB1 alleles in 23 unrelated individuals (Supplementary Table 3). Among the HLA class I alleles, we noted the highest frequencies for A*02:01 (∼63%), B*27:05 (∼23%) and C*02:02 (∼17%). For the HLA class II alleles, we obtained the highest frequencies for DPB1*02:01 (∼37%), DQB1*03:01 (∼41%) and DRB1*08:01 (∼28%).

We noted 28 different mitochondrial DNA (mtDNA) and 9 Y-chromosome haplotypes (Supplementary Table 1). Interestingly, 9 of 25 males carried the same Y-chromosome haplotype.

We performed kinship analyses using f3 statistics^22^ and READ^34^. Both programmes identified one triplet consisting of a female and two males as first-degree relatives (Supplementary Fig. 5). Parent-child relationships can be ruled out as all three individuals died in infancy (at age 1-3 years) or early childhood (4-6 years). This leaves the sibling constellation as the only other possible explanation, which is supported by the respective mtDNA and Y-chromosome haplotypes as well as HLA allele profiles.

## Discussion

It has clearly been established that the transformation from the LBK, which is characterised by a homogeneous material culture over a large area, to the later more diverse Neolithic cultures in Europe was accompanied by turnovers in the genomic record^3^. However, the population interactions underlying this transformation have not yet been fully analysed. The admixture events were geographically highly localized and involved various populations with different ancestry components^3^. These processes likely led to the increase in HG ancestry proportions and mtDNA lineages that were observed in Middle to Late Neolithic communities^1,7^. It is currently not known what might have influenced these wide-spread demographic and genomic processes in Europe, but climate change and/or social processes may be considered contributing factors^34^.

Here, we investigated a community of 42 Late Neolithic farmers excavated from the WBC gallery grave in Niedertiefenbach, Germany^16,17,36^. As expected, the studied population exhibited a mixture of genomic components from western HG and early farmers. The continuous range (34-58%) of the relatively high genetic HG proportion in the Niedertiefenbach collective is surprising. Admixture dating indicated that the mixing of the two components started 14.85 +/-2.82 generations before 3,300-3,200 cal. BCE. From these results, it cannot be inferred to what extent the contributing populations themselves were already admixed or which subsistence economy they practised. But interestingly, the estimated admixture date of 3,770-3,600 cal. BCE coincided with farming expansion phases and social changes during the Late MC (3,800-3,500 BCE)^37^. Archaeologically, there is a well-documented continuity from Late MC to WBC^9^. MtDNA data from two MC sites in France^38^ and Germany^39^ indicate that the analysed individuals belonged to an already admixed population comprising haplotypes typical of both farmers and HG^38^. Human genomic datasets from clear archaeological MC contexts are not available as yet. A possible exception could be the genome-wide data of four individuals from the Blätterhöhle that may be chronologically (based on their radiocarbon dates of 4,100-3,000 BCE) and geographically linked with Late MC and/or WBC^6^. However, it has to be kept in mind that the remains were found in a cave without any definite cultural assignment. Our analyses showed that the Niedertiefenbach population appeared most closely related to the Blätterhöhle collective. In particular, their large HG components (39-72%)^3^ fell into the range observed for Niedertiefenbach. Moreover, they were the best proxies for the HG and farmer components of the Niedertiefenbach sample. In addition, our ALDER admixture date is very similar to the one obtained for the Blätterhöhle that yielded 18-23 generations before the average sample date of 3,414 +/-84 cal. BCE^3^. Thus, there is a probable genetic link between the people buried in the Blätterhöhle and those in the gallery grave of Niedertiefenbach.

The WBC-associated population in Niedertiefenbach represents a genetically diverse group with a very broad spectrum in their HG proportions. This finding suggests that the admixture was still in progress at that time or had taken place a few generations before. This scenario is tentatively supported by the ALDER analysis (admitting that its admixture date may be biased towards a more recent time point). Given the surprisingly large HG component, it seems conceivable that the admixture included also individuals who had an exclusive or near-exclusive genetic HG ancestry. Taking into account all available lines of evidence, we hypothesize that the increase in the HG component likely occurred during the consolidation of the MC and/or the beginning of the WBC and could have involved also direct gene-flow from unadmixed local western HG into expanding farming populations.

The genetic data of the Niedertiefenbach sample, along with information obtained from archaeological and osteological analyses, sheds light on the community that used this gallery grave. In total, the skeletal remains of a minimal number of 177 individuals were recovered from the 7 m^2^ site, reflecting a very high occupancy rate for a collective WBC burial^35^. The sex distribution in the sample, which was assessed based on diagnostic skull elements, was similar to those described for other prehistoric populations^40^. Regarding age, we did not observe a numerical deficit of children that is often recorded for Neolithic cemeteries in Germany^36,41,42^. Thus, it is likely that the skeletal population of Niedertiefenbach represented a demographic cross-section of the group that was associated with this gallery grave. The phenotype reconstruction revealed that the examined individuals had a predominantly dark complexion and were genetically not yet adapted to digest starch-rich food or lactose. These phenotypes have typically been described for HG and early farmers^3^.

Overall, the genomic data indicate that the gallery grave was mainly used by not closely related people who may have lived in various neighbouring locations. This observation is supported by the large number of mtDNA (28) and Y-chromosome (9) haplotypes. However, also directly related individuals were interred. In one case, we observed inhumations of first-degree relatives (Supplementary Fig. 4). In addition, the presence of one frequent Y-chromosome haplotype suggests a patrilineage.

In line with studies investigating the health status of Neolithic populations in central Europe^43^, the Niedertiefenbach individuals showed numerous unspecific skeletal lesions that could be indicative of physical stress, including malnutrition, and infections^36^. Interestingly, we did not observe any pathogens. This observation is consistent with aDNA-based findings describing only relatively few sporadic cases of infectious diseases for the Neolithic period^44^. Noteworthy is the absence of *Yersinia pestis*, as lineages of this bacterium have already been postulated for the Late Neolithic and are reported in a Scandinavian case dated to 2900 cal. BCE^45^. If it were present, we would have expected to detect the pathogen, given the good preservation of endogenous DNA in the samples.

The HLA class I and II dataset generated for Niedertiefenbach was relatively small and thus precluded thorough statistical analysis. However, relative to contemporary European populations some striking shifts in allele frequencies could be seen (Supplementary Table 3). Interestingly, the majority of these alleles (e.g. B*51:01, DQB1*03:01) are today associated with higher resistance to viral pathogens (e.g. HIV, HCV, influenza A) and higher susceptibility to bacterial infections or complications thereof^46,47,48,49^. This observation strengthens the hypothesis that ancient epidemics influenced the present-day frequency of variants associated with modern inflammatory diseases^46,47^. Later on it may have lost its relative fitness advantage, for example because pathogens adapted to this most common allele in a process of negative frequency-dependent selection^50^, and was replaced by alleles beneficial against newly emerging human pathogenic bacteria, such as *Y. pestis*.

Further, notable difference concerns the HLA allele DRB1*15:01. It is widespread in present-day Europeans (ca. 15%), but absent in Niedertiefenbach samples. This allele predisposes to mycobacterial infections (tuberculosis, leprosy)^51^. In disease studies, the SNP allele rs3135388-T is often used as a marker for DRB1*15:01^52^. In the published aDNA datasets^25^, rs3135388-T was also found to be absent in all European Palaeolithic, Mesolithic and Neolithic populations analysed. It seemed to appear for the first time only during the Bronze Age. Since then, its initially high frequency (approx. 20%) has decreased to the present levels (Supplementary Fig. 6). This finding raises the intriguing possibility that the allele might have been incorporated into the European gene pool as part of the steppe-related ancestry component in the Final Neolithic and Bronze Age.

The advent of farming and subsequent shifts in pathogen exposure are thought to have radically changed the immune genes in early agriculturalists^25^. The immune response of the Niedertiefenbach collective was primarily geared towards fighting viral agents. To what extent this antiviral profile was due to the specific demographic history of the Niedertiefenbach population or was typical of Neolithic communities in general remains to be clarified. Together, our study detected large differences in the HLA variation and immune responsiveness over the last 5300 years in Europe.

By applying a comprehensive genomics approach to individuals interred in the WBC-associated collective burial in Niedertiefenbach, we discovered that the community, which used this site for about 100 years, was genetically heterogeneous and carried both Neolithic and HG ancestry components. The mixture of these two components likely occurred at the beginning of the 4^th^ millennium, indicating important demographic and cultural transformations during that time in western Europe. This event may also have affected the immune status of the admixed population and its descendants for generations to come.

## Methods

### Samples

The archaeological site and anthropological characteristics are described elsewhere^16,17,36^.

### Radiocarbon dating

Collagen was dated from 25 human bone samples, originally collected for aDNA analysis, and each attributed to a different individual. Dating was performed following standard protocols at the Leibniz Laboratory for AMS Dating and Isotope Research, Kiel (details in Supplementary Information).

### aDNA extraction and sequencing

Surface contaminations from petrous bones and teeth were removed with bleach solution. Partial uracil-DNA-glycosylase treated sequencing libraries were prepared from bone powder-derived DNA extracts following previously established protocols^29^. Sample-specific index combinations were added to the sequencing libraries^53^. Sampling, DNA extraction and the preparation of sequencing libraries were performed in clean-room facilities of the Ancient DNA Laboratory in Kiel. Negative controls were taken along for the DNA extraction and library generation steps. The libraries were paired-end sequenced using 2×75 cycles on an Illumina HiSeq 4000. Demultiplexing was performed by sorting all the sequences according to their index combinations. Illumina sequencing adapters were removed and paired-end reads were merged if they overlapped by at least 11bp. Merged reads were filtered for a minimum length of 30 bp.

### Pathogen screening

All samples were screened with MEGAN^54^ and the alignment tool MALT^19^ for their metagenomic content using parameters as described in Krause-Kyora et al. 2018^29^.

### Mapping and aDNA damage patterns

Sequences were mapped to the human genome build hg19 (International Human Genome Sequencing Consortium, 2001) using BWA 0.7.12^55^ with a reduced mapping stringency parameter “-n 0.01” to account for mismatches in aDNA. Duplicates were removed. C to T misincorporation frequencies were obtained using mapDamage 2.0^55^ in order to assess the authenticity of the aDNA fragments^18^. After the validation of terminal damage, the first two positions from the 5’ end of the fastq-reads were trimmed off.

### Genotyping

Alleles were drawn at random from each of the 1,233,013 SNP positions^1,2,25^ in a pseudo-haploid manner using a custom script as described in Lamnidis et al. 2018^57^. Datasets were filtered for at least 10,000 SNPs to be considered for further analysis^5^.

### Genetic sex determination

Sexes were determined based on the ratio of sequences aligning to the X and Y chromosomes compared to the autosomes^58^. Females are expected to have a ratio of 1 on the X chromosome and 0 on the Y chromosome, whereas males are expected to have both X and Y ratios of 0.5.

### Contamination estimation and authentication

Estimation of DNA contamination was performed on the mitochondrial level using the software Schmutzi^59^, and in males additionally on the X-chromosomal level by applying ANGSD^60^ to investigate the amount of heterozygosity on the X chromosome.

### Principal component analysis

The genotype data of the Niedertiefenbach collective was merged with previously published genotypes of 5519 ancient and modern individuals genotyped on the aforementioned 1,233,013 SNPs using the program *mergeit* from the *EIGENSOFT* package^61^. Principal component analysis (PCA) was performed using the software *smartpca*^61^ projecting the genotype datasets of the Niedertiefenbach and all other ancient individuals on the principal components calculated from genotype datasets of 59 West Eurasian populations by use of the ‘lsqproject’ option.

### ADMIXTURE analysis

Prior to ADMIXTURE analysis, we used Plink (v1.90b3.29) to filter out SNPs with insufficient coverage (0.999) and a minor allele frequency (maf) below 5%. LD pruning was performed to filter out SNPs at an R^2^ threshold of 0.4 using a window size of 200 and a step size of 25. We ran ADMIXTURE (version 1.3.0)^20^ on the same populations as used in the PCA analysis and a number of ancestral components ranging from 4 to 12. Cross-validation was performed for every admixture model.

### Admixture dating

The source code of ALDER (v1.03)^23^ was modified to decrease the minimal number of samples needed for the analysis, following a suggestion from this work: https://www.diva-portal.org/smash/get/diva2:945151/FULLTEXT01.pdf. In so doing also reference populations with only a single individual could be included.

The following reference populations were used for Niedertiefenbach: Anatolia_Neolithic, OrienteC_HG, Croatia_Mesolithic_HG, Bichon, Blatterhohle_MN, Koros_Hungary_EN_HG, Serbia_HG, Serbia_Mesolithic_Neolithic, Narva_LT, Iron_Gates_HG, Loschbour, Iberia_HG, Latvia_EN, Baalberge_MN France_MN, Latvia_HG.

To calculate calender dates of admixture we multiplied the obtained the average ALDER generation time for two-way admixture models with significant LD-decay curves with an assumed generation time of 29 years^24^.

### F3 outgroup statistics

f3 outgroup statistics were run as a part of the *Admixtools* package^22^ in the form of *f*_*3*_(*Niedertiefenbach; test, Mbuti*) using for *test* the same populations as in the PCA and ADMIXTURE analysis.

### qpADM analysis

qpADM analysis was run on transition-filtered genotypes that were previously prepared for ADMIXTURE analysis as described above. We ran 48 different combination models of Niedertiefenbach as a two-way admixture, since three-way admixture models appeared to be less feasible indicating that the 3^rd^ component was excessive. The following populations were used as outgroups: Mbuti, Ust Ishim, Kostenki14, MA1, Han, Papuan, Onge, Chukchi and Karitiana.

### Kinship analysis

Kin relatedness was assessed using READ^34^ and lcMLkin^62^. READ identifies relatives based on the proportion of non-matching alleles. lcMLkin infers individual kinship from calculated genotype likelihoods. A pair of individuals was regarded related only if evidence of relatedness was independently provided by both programs (Supplementary Fig. 5).

### Determination of mitochondrial and Y chromosome haplotypes

Sequencing reads were mapped to the human mitochondrial genome sequence rCRS^63^. Consensus sequences were generated in Geneious (v. 9.1.3) using a default threshold of 85% identity among the covered positions and a minimum coverage of 3. HAPLOFIND^64^ was applied to assess mitochondrial haplotypes from the consensus sequences and yHaplo^65^ to determine Y chromosome haplotypes in male individuals.

### Calling of phenotypic SNPs

We generated a pile-up of reads mapping to the positions of the selected phenotypic SNPs with samtools mpileup (v. 1.3) in order to see how many reads supported which allele for each individual.

### HLA typing and analysis

We used a previously established HLA capture and HLA typing pipeline^29^. In addition, we applied OptiType^30^ for automated HLA class I and II typing. We then removed one in a pair of datasets of directly related individuals (1^st^ and 2^nd^ degree relatives) based on the maximum number of reads supporting the HLA call in either of the related individuals. Samples with low coverage of the HLA region were also excluded. Only alleles that were consistently called by both methods were considered for the analysis. For comparing the ancient HLA allele pool with a representative modern allele pool, we used a cohort of 3,219 healthy northern German individuals and imputed HLA genotypes at 2^nd^ field level of resolution from high-density SNP data following an established procedure^66^.

## Data availability

The aligned sequences are available through the European Nucleotide Archive under accession number XXXXXXXXX.

## Acknowledgements

This study was funded by the Deutsche Forschungsgemeinschaft (DFG, German Research Foundation) through Projektnummer 2901391021 SFB 1266 to B.K.-K. and A.N., grant LE 2593/3-1 to T.L.L. and Germany’s Excellence Strategy – EXC 2167-390884018. AN was supported by the Dorothea Erxleben Female Investigator Award of the DFG Cluster of Excellence Inflammation at Interfaces (EXC306). We are grateful to Stephan Schiffels, Thiseas C. Lamnidis and Alissa Mittnik from the Max Planck Institute for the Science of Human History in Jena for sharing their expertise in ALDER, qpADM and kinship analyses and their advice on result verification. We acknowledge financial support by Land Schleswig-Holstein within the funding programme Open Access Publikationsfonds. J.B. and F.P. were funded by the International Max Planck Research School for Evolutionary Biology.

## Author contributions

B.K.-K., A.N. and Ch.R. conceived and designed the research. K.F. analyzed the human skeletal remains. L.B. and B.K.-K. generated ancient DNA data. A.I., J.S. and B.K. analyzed the ancient DNA data. A.I., A.S., F.P., A.F., L.B., J.D., J.B., D.E., J.Ch.K., R.B., O.K., I.M., T.L., A.F., J.K. and B.K.-K. analyzed modern and ancient HLA data. S.Sch., J.M., Jo.M., Ch.R., C.D., M.F., A.N. and B.K.-K. interpreted the findings. A.N., A.I. and B.K.-K. wrote the manuscript with input from all other authors.

